# Fruit performance of kiwifruit in the loess area of Northwest China

**DOI:** 10.1101/633206

**Authors:** Kaibin Guo, Zhen Guo, Yun Guo, Guang Qiao

## Abstract

Attempts had been made to provide evident insight into the performance of fruit quality and the status of soil property, and explicate the soil property factors that dominantly affect the fruit quality of kiwifruit. Currently, 8-year-old kiwifruit cultivar ‘Hayward’, which was grown in Zhouzhi County (108°37′ E, 33°42′N), Shanxi Province of China, was used as materials. The results of Pearson correlation coefficient illustrated that the soil organic matter (SOM) content was positively related to soil properties except the soil PH. Moreover, based on the canonical correlation analysis (CCA), canonical variables alkaline hydrolyzable-N (AN), available ferrum (AFe), available boron (AB), PH in soil property index and the fresh weight of single fruit (FW), fruit shape index (FI), total soluble solids (TSS), titratable acidity (TA), total soluble sugar (SS) in fruit quality parameter were selected. And the ‘best’ regression equation (model) indicated that the effects of soil property somewhat varied among ‘Hayward’ fruit qualities in the loess area of Northwest China. Specifically, FW and SS could be mainly affected by soil AN, and FI affected by soil AB and PH. Fruit SS mostly depended upon soil AFe, whereas TSS was affected by soil AN, AFe and PH. The effect of soil PH on fruit quality is probably achieved, however, affecting the absorption of soil nutrients.

## Introduction

As an edible berry of several *Actinidia* species, kiwifruit is unequalled, compared with other commonly consumed fruit, for their rich antioxidant, health benefits, and consumer appeal [1]. The successful development of kiwifruit industry was first appeared in New Zealand, and then expanded rapidly in other countries. Over the past decade, large amount of kiwifruit had been planted in China due to the strongly economic incentive caused by the international kiwifruit trade [2]. Among many factors that affect economic benefit of fruit, fruit quality plays a pivotal role in consumption choice, agro-industrial processing, food technology, and production data.

Normally, the fruit quality was determined by both cultivar and growing environment, which was confirmed on many tree species, *e.g.* blueberry [3], citrus [4], apple [5]. Likewise, the difference on fruit quality of kiwifruit in terms of sensory acceptance, sugar and nutritive factors as well as polyphenols among cultivars were reported by previous studies [6,7], which was affected by tree canopy density [8], ambient temperature [9], soil water potential as well [10]. Particularly, previous surveys illustrated that soil nutrient is of utmost importance for fruit quality of kiwifruit. For instance, Hasinurrahman et al. [11] provided a indirect evidence that kiwifruit quality mostly depended upon the use of a biological farming system, which linked with the soil properties. Lago et al. [12] found that the K: Mg ratio in soil was closely related to the absorption of Ca^2+^ and Mg^2+^ by kiwifruit. The level of soil nutrient, however, majorly depended on the environmental conditions, *e.g*.,topography, climate and man-induced factors, *e.g*., agricultural management [13,14]. For instance, Khan et al. [4] reported that the orchard location and soil nutrient status play a crucial role in fruit quality of citrus in Sargodha city of Pakistan. Zhouzhi, a county of Guanzhong plain, Shanxi Province of China, where the soil type are primarily composed of anthrosol, which is super common type in the loess area of Northwest China. Researchers had previously applied the Pearson correlation coefficient for predicting the relationship between fruit quality of kiwifruit and anthrosol type soil in Northwest China. But in the present work, the canonical correlation analysis (CCA) method was applied to select the main soil properties factors affecting fruit quality. And then the method of stepwise multiple regression was used for establishing the ‘best’ regression equation (model). Therefore, the objectives of the work are: (a) assessing of the performance of fruit quality and the status of soil nutrient, and exposing of the mutual relationship; (b) based on the method of multivariate analysis, illustrating the soil nutrient factors that dominantly affect the fruit quality of kiwifruit in Northwest China.

## Materials and methods

### Plant material and soil sample

Currently, a well-known kiwifruit cultivar ‘Hayward’, which was grown in Zhouzhi County (108°37′E, 33°42′N), Shanxi Province of China (Fig 1), were sampled as materials in 2018. The climate of the orchard belongs to a warm temperate continental monsoon climate, characterized by the average total annual precipitation of 850.52 mm and the mean annual sunshine hours of 1,774.6 h as well as average annual temperatures of 14.3 °C. The 8-year-old kiwifruit trees were planted in the row spacing of 4 m×5 m and grown to the height about 4.5 m. At the stage of fruit mature (from early August to end September), a sample of randomly obtained 12 fruits were harvested from East, South, West and North of each tree (three trees per replication), and quickly frozen in ice box with under −20 °C. Soil in this area is mainly a type of earth-cumuli-orthic anthrosols. The average soil bulk density in soil layer depth 0-40 cm is about 1.35-1.62 g·cm^−3^. To approach more representative data, soil was sampled randomly from three distantly separated plots, which was consisted of three replicates of nine trees, from June to August. Soil in 0-40 cm layer was collected with soil drilling method from East, South, West and North of each tree crown vertically inward 50 cm, and were air-dried naturally at room temperature and crushed, before been sieving through a 2-mm mesh sieve.

**Fig 1.**
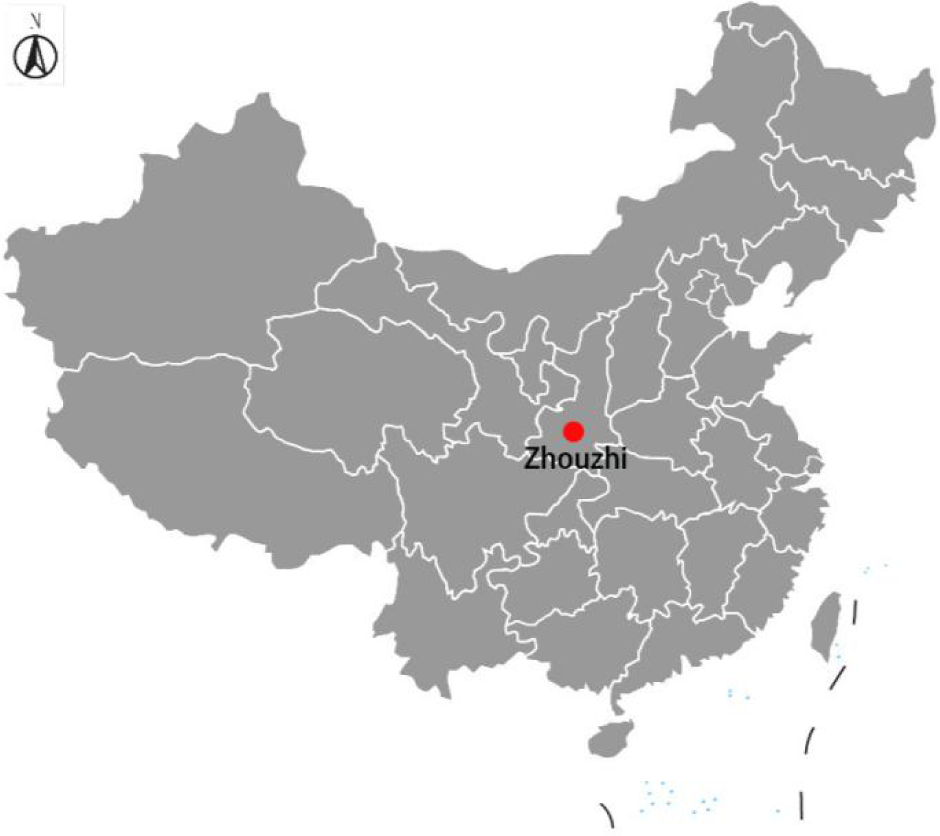
Geographical location of sampling sites in China.

### Determination the performance of fruit quality

#### Weight and shape index

The fresh weight of single fruit (FW, expressed in g) was determining on an analytical balance (Mettler-Toledo AL204). The fruit height and width (expressed in mm) were measuring by a Vernier caliper (Nscing Es, China) and calculating fruit shape index (FI) as height/width ratio. Three replications were conducted and averaged the value.

#### Soluble solids, titratable acidity, soluble sugar

These levels were assessed with juice obtained randomly from three fruits per replication. Juice was obtained from fruit center using the indelicate super automatic extractor (SamTech, Mannersdorf, Austria). The concentration of total soluble solids (TSS) were determined at 20 °C using a refractometer (SHGWYT-2, 0–32%, China) and reported as °Brix. Titratable acidity (TA) was determined by titration with 0.05 mol L^−1^ NaOH to PH 8.2 and reported as g citric acid per 100 g fresh weight. Total soluble sugar (SS) was measured through anthrone-H_2_SO4 colorimetry and expressed in °Brix.

#### Detection the property of soil

Total organic matter content was gravimetrically determined by loss on ignition (OM_LOI_). Briefly, 10 g samples were oven-dried at 105 °C overnight, weighed, ignited to equilibrium in a muffle furnace set at 350 °C for 18 h and reweighed. The lower ignition temperature was chosen to prevent errors from high loss of structural water from kaolinite clays which is generally greatest between 450 and 600 °C [15]. The total nitrogen (TN) was measured by the Kjeldal method, while PH was determined using 1:2.5 CaCl_2_ dilution method [16]. Alkaline hydrolyzable-N (AN) in soil was assessed with a micro-diffusion technique after samples went through alkaline hydrolysis. Soil available phosphorus (AP) was determined by the Olsen method [17]. Soil available potassium (AK) was measured in 1 mol L^−1^ NH_4_ OAc extracts by flame photometry. DTPA-extractable micronutrients, *i.e*., available ferrum (AFe), available Zinc (AZn), available boron (AB) contents in soil were measured on atomic absorption spectrophotometer (4530F China) from 1:2 soil-extractant ratio using DTPA-TEA buffer (0.005 *M* DTPA + 0.001 *M* CaCl_2_ + 0.1 *M* TEA, pH 7.3) with method described by Lindsay & Norvell (1978) [18].

#### Statistical Analysis

Pearson’s correlation analysis (PCA) was used to evaluate the Pearson correlation coefficient between the fruit quality values and soil nutrient levels by SPSS 22.0 statistical software. And heatmap was drawn by R package function heatmap.2 (http://cran.r-project.org/web/packages/gplots/index.tml).

According to the modern regression theory, soil nutrients and fruit qualities belong to two different normal distributions, and each variable set consisting of more than two variables. To model the response variable as a function of one or more predictor variables. CCA completing by ‘Sytax’ macro program of SPSS, was employed for assessing the relationship between the soil nutrients, *i.e*., OM(x1), TN(x_2_), AN(x_3_), AP(x_4_), AK(x_5_), AFe(x_6_), AZn(x_7_), AB(x_8_), PH(x_9_) as independent set of variables and fruit qualities, *i.e*., FW(y_1_), FI(y_2_), TSS(y_3_), SS(y_4_), TA(y_5_) as dependent set. Then, the method of stepwise multiple linear regression (SMLR) was used alternatively to build linear regression prediction equations (models) by SPSS 22.0 software.

## Results

### Quality performance of fruit and property status of soil

Performance of fruit quality were presented in Table 1. The FW values ranged from 77.63 to 91.12 g with a mean of 84.49 g (STD=5.12, *C.V.* =6.00%), while FI levels varied from 1.09 to 1.30 with an average of 1.20 (STD=0.07, *C.V.* =6.00%). As for the internal qualities of kiwifruit fruit, *i.e*., TSS, SS, and TA, which were used to calculate values of the TSS/TA and SS/TA. Contents of TSS and SS ranged from 12.99 to 14.29 °Brix with a mean of 13.62 °Brix (STD=0.58, *C.V.* =4.00%) and 8.57 to 10.00 °Brix with an average of 9.01 °Brix (STD=0.66, *C.V.*=7.00%), respectively. Whereas TA concentration varied from 1.12 to 1.33% with a mean of 1.23% (STD=0.08, *C.V*.=7.00%). The TSS/TA and SS/TA estimates varied from 9.77 to 11.95 with a mean of 11.16 (STD=0.89, *C.V*. =8.00%) and 6.11 to 8.93 (STD=0.98, *C.V.* =13.00%).

**Table 1.** Fruit quality of ‘Hayward’ kiwifruit.

|  | FW | FI | TSS | SS | TA | TSS/TA | SS/TA |
| --- | --- | --- | --- | --- | --- | --- | --- |
|  | (g) |  | (°Brix) | (°Brix) | (%) |  |  |
| Maximum | 77.63 | 1.30 | 14.29 | 10.00 | 1.33 | 11.95 | 8.93 |
| Minimum | 91.12 | 1.09 | 12.99 | 8.57 | 1.12 | 9.77 | 6.11 |
| Mean | 84.49 | 1.20 | 13.62 | 9.01 | 1.23 | 11.16 | 7.39 |
| STD | 5.12 | 0.07 | 0.58 | 0.66 | 0.08 | 0.89 | 0.98 |
| *C.V.* (%) | 6.00 | 6.00 | 4.00 | 7.00 | 7.00 | 8.00 | 13.00 |

Note: ‘STD’ and ‘C.V.’ stand for standard deviation and coefficient of variation respectively.

Status of the studied soil properties were shown in Table 2. The OM contents of the samples varied from 22.75 to 24.10 g·Kg^−1^ with an average of 23.43 g·Kg^−1^ (STD=0.53, *C.V.* =2.27%). TN and AN statues with different order of magnitude, ranged from 1.46 to 1.61 g·Kg^−1^ with a mean of 1.55 g·Kg^−1^ (STD=0.06, *C*.*V*. =3.81%) and 104.95 to 109.05 mg·Kg^−1^ with an average of 107.17 mg·Kg^−1^ (STD=1.76, *C*.*V.*=1.64%) respectively. Moreover, the estimates of AP, AK, AFe with an average of 94.45 mg·Kg^−1^, 110.95 mg·Kg^−1^, 9.11 mg·Kg^−1^, varied from 89.86 to 106.73 mg·Kg^−1^(STD=6.37, *C.V.* =6.74%), 107.32 to 114.01 mg·Kg^−1^ (STD=2.83, *C.V.*=2.55%) and 8.73 and 9.51 mg·Kg^−1^ (STD=0.27, *C.V.* =2.91%), respectively. Furthermore, the AZn and AB values of sampled soils ranged from 0.92 to1.08 mg·Kg^−1^ with a mean of 0.98 mg·Kg^−1^ and 0.53 to 0.63 mg·Kg^−1^ with an average of 0.57 mg·Kg^−1^. Afterwards, the levels of soils PH varied from 6.15 to 7.39 with an average of 6.95 (STD=0.42, *C.V*. =6.06%).

**Table 2.** Nutrient status of the studied soil.

|  | OM | TN | AN | AP | AK | AF<sub>e</sub> | AZn | AB | PH |
| --- | --- | --- | --- | --- | --- | --- | --- | --- | --- |
|  | g·Kg<sup>-1</sup> | g·Kg<sup>-1</sup> | mg·g<sup>-1</sup> | mg·Kg<sup>-1</sup> | mg·Kg<sup>-1</sup> | mg·Kg<sup>-1</sup> | mg·Kg<sup>-1</sup> | mg·Kg<sup>-1</sup> |  |
| Maximum | 24.10 | 1.61 | 109.05 | 106.73 | 114.01 | 9.51 | 1.08 | 0.63 | 7.39 |
| Minimum | 22.75 | 1.46 | 104.95 | 89.86 | 107.32 | 8.73 | 0.92 | 0.53 | 6.15 |
| Mean | 23.43 | 1.55 | 107.17 | 94.45 | 110.95 | 9.11 | 0.98 | 0.57 | 6.95 |
| STD | 0.53 | 0.06 | 1.76 | 6.37 | 2.83 | 0.27 | 0.06 | 0.04 | 0.42 |
| *C.V.* (%) | 2.27 | 3.81 | 1.64 | 6.74 | 2.55 | 2.91 | 6.19 | 7.61 | 6.06 |

Results of correlation analyses of fruit qualities and soil properties were shown in Fig 2. As for the soil properties, OM content showed a positive correlation with others measured parameters except for the soil PH, which showed significant negative correlation with AP level (*R*=−0.860*). Moreover, obvious correlation among some fruit traits were found as well, such as FW was positively and significantly correlated with SS content (*R*=0.813*) and TSS/TA (*R*= 0.885*); SS content was significantly positive correlated with TSS/TA (*R*= 0.889*) and SS/TA (*R*= 0.964**), whereas remarkably negative correlation with TA concentration (*R*=−0.821*); TSS/TA showed significantly positive correlation with SS/TA (*R*= 0.898*) while obviously negative correlation TA (*R*=−0.862*), which showed significantly negative correlation with SS/TA (*R*=−0.940**). Furthermore, statistically significant positive or negative correlation between fruit traits and soil properties were observed. Soil AN content was negatively and significantly correlated with fruit TA (*R*=−.955**), however, positively and obviously correlated with TSS/TA (*R*=0.863*) and SS/TA (*R*=0.902*). Soil Fe content showed significantly positive correlation with fruit TSS concentration (*R*=0.838*).

**Fig 2.**
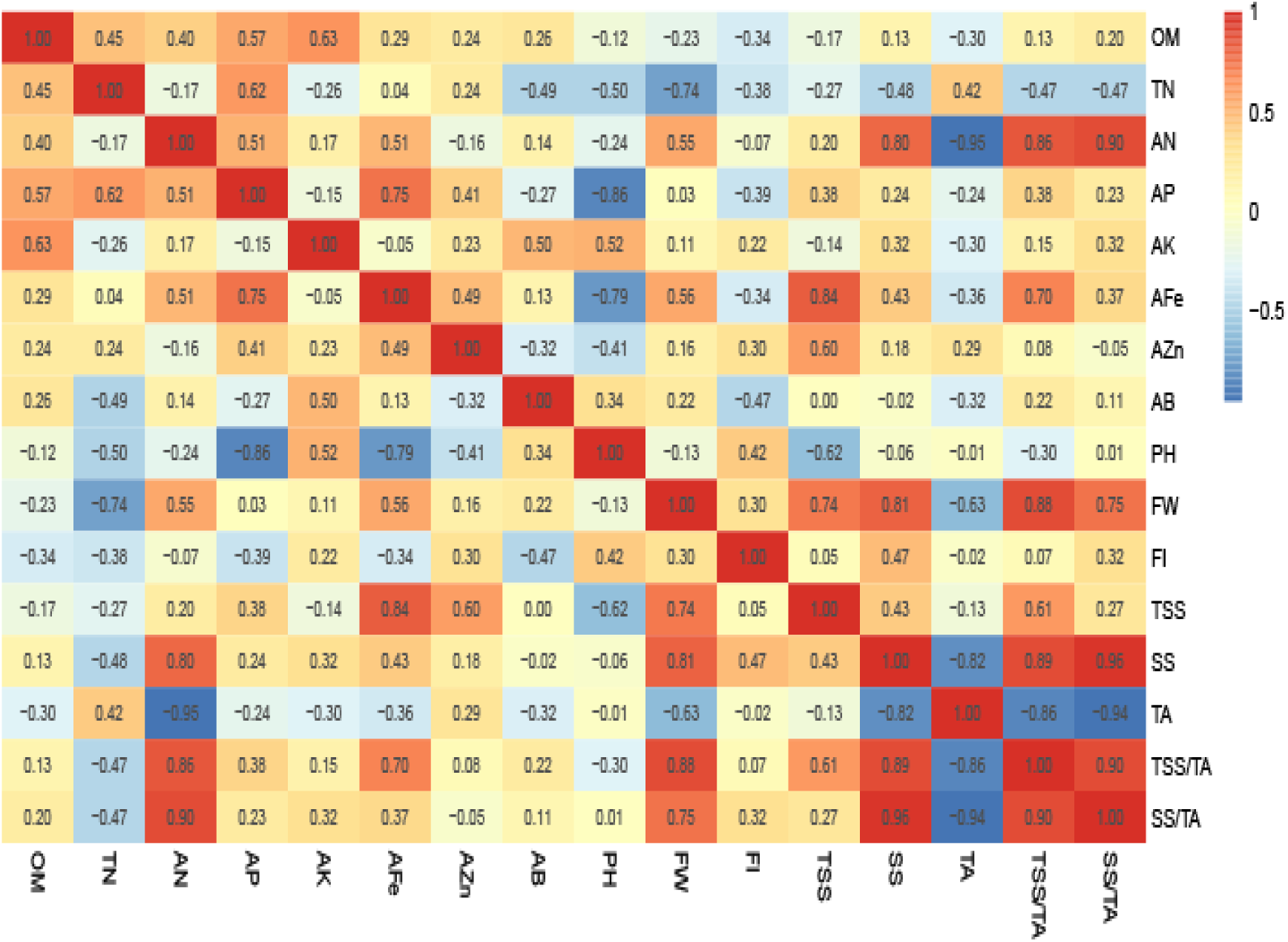
Pearson correlation heatmap of fruit qualities and soil nutrients.

### Multivariate analysis of relationship between soil property and fruit quality

#### CCA of soil nutrient and fruit quality

Results of CCA were revealed in Table 3. According to Wilk’s λ and Chi-square tests for canonical correlation coefficients, three pairs of canonical variables (*U*_*1*_,*V*_*1*_; *U*_*2*_,*V*_*2*_; *U*_*3*_,*V*_*3*_) of soil property and fruit quality, of which the canonical correlation coefficient and *p*-value for these canonical variables were 1.000 and 0.000, 1.000 and 0.000, 0.987 and 0.001 respectively, all reached a significant level. The three pairs of canonical variables for soil property and quality could be described as follows:

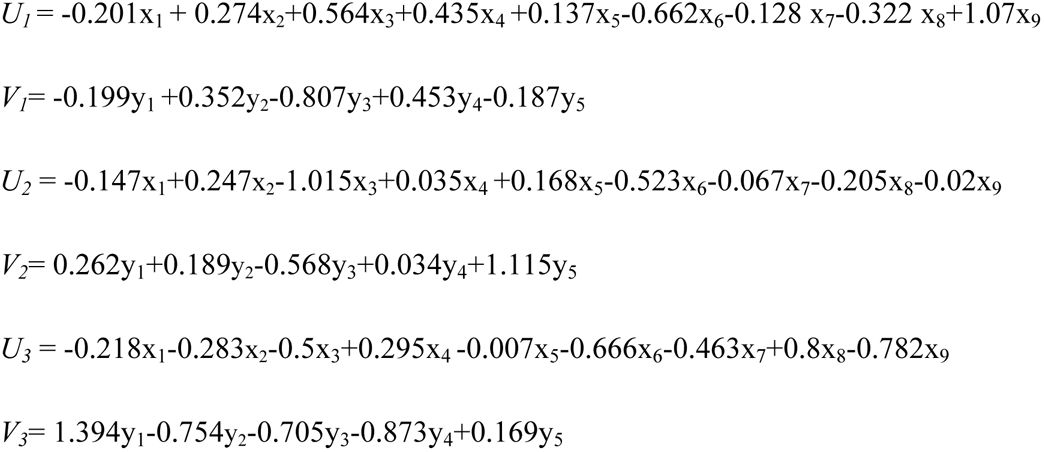

**Table 3.** Wilk’s λ and Chi-square tests of canonical correlation coefficients.

| canonical<br>vector No. | canonical correlation<br>coefficients | Wilk's $\lambda$ | $\chi^2$ | df | $p$ |
| --- | --- | --- | --- | --- | --- |
| 1 | 1.000 | 0.000 | 869.829 | 45 | 0.000** |
| 2 | 1.000 | 0.000 | 463.298 | 32 | 0.000** |
| 3 | 0.987 | 0.005 | 61.430 | 21 | 0.001** |
| 4 | 0.894 | 0.188 | 19.246 | 12 | 0.083 |
| 5 | 0.253 | 0.936 | 0.763 | 5 | 0.979 |

Note: '\*\*' indicated significance at $p < 0.01$ level

**Table 4.** The stepwise regression equations and *F*-value for soil nutrient to fruit quality.

| Fruit qualities | Soil nutrients | Regression equations | $R^2$ | $F$-value |
| --- | --- | --- | --- | --- |
| FW($y_1$) | OM($x_1$), TN($x_2$) | $y_1 = 1.18x_3 - 41.878$ | 0.191 | 5.496* |
| FI($y_2$) | AN($x_3$), AP($x_4$), | $y_2 = -1.377x_8 + 0.09x_9 + 1.731$ | 0.366 | 6.475** |
| TSS($y_3$) | AK($x_5$), AFe($x_6$) | $y_3 = 0.884x_6 + 5.541$ | 0.525 | 21.999*** |
| SS($y_4$) | AZn($x_7$), AB($x_8$) | $y_4 = 0.188x_3 - 11.124$ | 0.258 | 7.610* |
| TA($y_5$) | PH($x_9$) | $y_5 = -0.045x_3 - 0.028x_6 - 0.072x_9 + 6.771$ | 0.971 | 215.303*** |

Note: '\*', '\*\*' and '\*\*\*' indicated significance at $p < 0.05$ , $< 0.01$ , $< 0.001$ level, respectively.

For the first pair of canonical variables, AN(x_3_), AFe(x_6_), PH(x_9_) in soil nutrient index and TSS(y_3_) in fruit quality index presented greater weight coefficient. It illustrated that the contents of soil AN, AFe and PH value played a crucial role in the levels of fruit TSS. The second pair of canonical variables (*U*_*2*_,*V*_*2*_) where the dominant factors are AN(x_3_), AFe(x_6_) in soil nutrient index, and TSS(y_3_), TA(y_5_) in fruit quality index. Similarly, the results indicated that the contents of AN, AF have dramatic impact not only on fruit TSS level, but also on TA concentration. Furthermore, AFe(x_6_), AB(x_8_), PH(x_9_) in soil nutrient index and FW(y_1_), FI(y_2_), TSS(y_3_), SS(y_4_) in fruit quality index of the third pair of canonical variables appeared a larger weight coefficient as well. It also showed that FW, FI, TSS, SS were obviously affected by the contents of soil AFe, AB, and PH value. Additionally, with canonical redundancy analysis, 9.1%, 12.9%, and 11.3% of the variance of soil nutrient could be explained by the canonical variables *V*_*1*_, *V*_*2*_, and *V*_*3*_, respectively. And 21.9%, 27.6%, and 14.8% of the variance of fruit quality could be explained by the canonical variables *U*_*1*_, *U*_*2*_, and *U*_*3*_.

#### Stepwise regression analysis for the nutrient to fruit quality

By establishing SMLR equation between dependent and independent variable, and automatically eliminate from the selected equation the insignificant independent variables, and *F-*value were used to evaluate statisical significance of SMLR model. The results of SMLR analysis indicated that AN(x_3_) were positively related to FW (y_1_) and SS(y_4_), whereas was negatively related to TA(y_5_). Moreover, AFe(x_6_) showed positive correlation with TSS(y_3_) while negative correlation with TA(y_5_). Furthermore, AB(x_8_) was negatively related to FI(y_2_). Additionally, PH(x_9_) was positively related to FI(y_2_), but negatively related to TA(y_5_).

## Discussion

Currently, difference in status of soil property was observed when comparing with the same soil type and land using same patial distribution, same climate condition in other regions of China [19–21]. It was elucidated that the level of soil property was affected by many interacting factors. Moreover, fruit quality of ‘Hayward’ cultivar in the studied orchard also has discrepancy to other orchards [22,23]. It was likely indicate that orchard location had significant effect on fruit quality, which was in accordance with the study by Shiri et al. [24], who revealed the significant difference in biochemical compounds of ‘Hayward’ kiwifruit fruit sampled from two different commercial “Hayward” orchards in the north of Iran. Furthermore, a growing tonnage of evidences have illustrated the closely relationship between soil property and fruit quality in many fruit species, *e.g*., in citrus [25], blueberry [3], apple [5]. Also, our finding explicated correlated relationship between soil nutrient and fruit quality of kiwifruit, among which soil AN has significantly negative correlation with fruit TA (*R*=−.955**) and AFe showed significant positive correlation with fruit TSS (*R*=0.838*). SS has significantly positive correlation with FW (*R*=0.813*). The level of SS in fruit closely correlated with photosynthetic capacity of tree, and the higher the ability can be conducive to the accumulation of dry matter and therefore enhancement of fruit mass, which was in accordance with the evidence from Bertin et al [26], who illustrated that the total photosynthetically active radiation (PAR) received during the production stage from flowering to harvest has obvious correlation with the fresh mass of tomato fruit. However, according to the report published by El-Gizawy et al. [27], increasing shading levels from 35% to 63% increased the TA in tomato fruit, it declared that the photosynthetic capacity has negative correlation with TA, which was consistent with our results that SS was remarkably and negatively correlated with TA (*R*=−0.821*).

It is well known that the synergistic and antagonistic effects existing among soil nutrients, which will affect the availability of soil nutrient, and therefore has an great impact on its uptake by crops [28,29]. Based on the CCA and SMLR, our investigation indicated that soil AN content has positive correlation with FW, which was in accordance with the evidence from Hipps et al [30], who explicated that the enhancement of nitrogen fertiliser application could enhance the FW of apple. Moreover, we found that soil AN and AFe content were positively related to the fruit SS and TSS concentration, respectively, which were in agreement with results of PCA. Lindsay [31] who previously reported that the activity of Fe in soil solution was highly PH-dependent. The significantly positive effect of soil AFe on fruit TA level was presented, in the current work, by a combination of CCA and SMLR. It is likely that soil AFe had an antagonistic effect against soil PH [32]. Soil PH, however, is one of the most important soil chemical properties in determining plant growth of nutrient uptake, and its effect on fruit quality was mainly achieved through the uptake of soil nutrients by plants [33,34].

## Conclusion

Our study provided useful information on the status of soil property in Guangzhong plain of Northwest China, and its effect on the performance of fruit quality of ‘Hayward’ cultivar. The content of SOM was positively related to the measured soil nutrients except for the soil PH. It was indicated that the effects of soil properties somewhat varied among ‘Hayward’ fruit qualities. Among these fruit quality performance, FW and SS mainly affected by soil AN, and FI affected by soil AB and PH. Fruit SS mostly depended upon soil AFe, whereas TSS was affected by soil AN, AFe and PH. The effects of soil PH on fruit quality is probably achieved, however, affecting the uptake of soil nutrients.

## Acknowledgments

The project was supported by grants from the High-level Innovation Talent Program of Guizhou Province, P. R. China.

## Author Contributions

Conceptualization: Kaibin Guo, Zhen Guo, Guang Qiao.

Data curation: Kaibin Guo, Zhen Guo.

Formal analysis: Yun Guo.

Funding acquisition: Yun Guo, Guang Qiao.

Investigation: Kaibin Guo, Zhen Guo, Guang Qiao.

Methodology: Guang Qiao.

Project administration: Yun Guo, Guang Qiao.

Resources: Guang Qiao.

Software: Kaibin Guo, Yun Guo.

Supervision: Guang Qiao.

Validation: Kaibin Guo.

Writing – original draft: Kaibin Guo.

Writing – review & editing: Kaibin Guo, Zhen Guo, Guang Qiao.

